# Static stability and swim bladder volume in the bluegill sunfish *Lepomis macrochirus*

**DOI:** 10.1101/2022.08.08.503202

**Authors:** Michael Fath, Stacy Nguyen, Joan Donahue, Sarah McMenamin, Eric Tytell

## Abstract

The static stability of a fish in the water is determined by the relative locations of its center of mass and center of buoyancy. These locations may not be constant, but may depend on the volume of the swim bladder. Changes in swim bladder volume affect both the distribution of mass and the overall volume of the fish, thus affecting the location of the center of mass and center of buoyancy, respectively. To determine the static stability and to examine the influence of the swim bladder on static stability, we used micro-computed tomography to estimate the locations of the center of mass and center of buoyancy in bluegill sunfish (*Lepomis macrochrius*). In fish oriented head up in the scanner, we found that the center of buoyancy is located 0.441 body lengths from the snout and 0.19 body lengths above the ventral surface of the pelvic girdle, and that the center of mass is 0.0012 BL posterior to and 0.00045 BL ventral to the center of buoyancy. Swim bladders from our specimens ranged in size from 1.9% to 7.6% total body volume, and we found no correlation between swim bladder volume and the distance between the center of mass and center of buoyancy. However, in fish scanned in a head-down orientation, the center of mass is anterior and dorsal to the center of buoyancy—the opposite configuration of that found in fish oriented head up. This change in center of mass and center of buoyancy seems to be caused by changes in the location of the swim bladder in the body: the centroid of the swim bladder is located more posteriorly in fish oriented head-down. The air in the bladder “rises” while heavier tissues “sink,” driving a change in tissue distribution and changing the location of the center of mass relative to the center of buoyancy. We conclude that while static stability does not change with swim bladder volume, pitch angle could have a marked effect on static stability.

## Introduction

In still water, any submerged object experiences two forces: gravity and buoyancy. The relative magnitude of these forces determines whether an object floats or sinks, and the balance of these forces on the object determines its stable orientation. To predict that orientation, the gravitational and buoyant forces can be treated as acting at distinct points in the object. The buoyant force acts upward on the center of buoyancy (COB), a point located at the geometric center (also called the centroid) of the displaced fluid volume (Smits, 2000). The gravitational force pulls downward at a location called the center of mass (COM), which can be estimated by dividing the object up into many small elements and taking the average location of all the elements, weighted by their mass (Walker, 2015). The location of the COM is thus determined by the distribution of mass and the COB is determined by the distribution of volume. Therefore, the two points can have different locations in heterogeneous objects. If the COM and COB are in different locations, there are exactly two equilibrium orientations: the COM directly above or directly below the COB. In both of these cases, the lines of action of the gravitational and buoyant forces are aligned and generate no torques. However, only the orientation with the COB above COM is stable; that is, small perturbations causing a misalignment of the two points will result in torques which act to move the COB back above the COM.

The statically stable orientation of a fish, like any other object, is one in which the COB is above the COM. However, unlike inanimate objects, fish also have a desired orientation (most often dorsal side-up) which may or may not also be the statically stable orientation of the fish. Indeed anecdotal and experimental evidence suggest that a horizontal, dorsal side-up orientation is not stable. Fish generally pitch or roll “belly up” when anesthetized or dead (Harris, 1936). Aleyev (1977) found horizontal displacements between the COM and COB in many fishes, indicating a pitching destabilization. Fish counter destabilizing torques with constant fin motions. This establishes dynamic stability: active body motions that maintain a desired orientation. More recently, Webb and Weihs (1994) explored hydrostatic stability in rock bass, perch, bluegill, and eel. Contrary to Aleyev’s earlier studies (1977), Webb and Weihs (1994) found no significant difference between the mean longitudinal locations of the COM and COB. Additionally, when they measured COM and COB locations using a plumb line technique, they found that in bluegill, the two points are statistically the same, suggesting that they are neutrally stable. They proposed that their apparently contradictory result was a result of the small distance between the COM and COB and the relatively larger variability they recorded from multiple measurements of each point.

Further complicating the relationship between the location of the COM and COB in fishes is the presence of the swim bladder. The swim bladder is a gas-filled organ which is dramatically less dense than the rest of the fish. The gas within the bladder compresses or expands as the fish changes depth. Fish can add or remove gas to the bladder in response (Steen, 1970) but this process is slow. Changes in swim bladder volume could affect the location of the COM by changing the distribution of mass throughout the body. These changes could also alter the location of the COB if the external shape of the fish changes: for example, if it swells in the region near the swim bladder.

In this study, we aimed to determine the locations of the center of mass and center of buoyancy as well as the relationship between swim bladder volume and their locations in the bluegill sunfish (*Lepomis machrochirus)*. We used micro-computed tomography (μCT) to calculate the location of the COM and COB in fish with untreated swim bladders and in fish with air removed from the swim bladder. In objects of uniform density, the COM and COB would be located in the same place. Non-uniform distribution of mass, as in fish, causes a displacement between these points. Therefore, we predicted that fish with relatively larger swim bladders would show a greater distance between the COM and COB due to the increased heterogeneity of tissue density.

## Methods

### Animal Preparation and Scanning

We collected 13 bluegill sunfish from White Pond in Concord, Massachusetts, in the summer of 2019. Prior to scanning, the fish were held in 10 gallon tanks. Fish were euthanized using an overdose of buffered MS-222 (Acros Organics) solution (0.04%) in 1 L water.

To generate a range of swim bladder volumes, we used a syringe to add or subtract air from the swim bladder. We estimated swim bladder volume using the relationship from McNabb and Meecham (1971) that the volume of the swim bladder is 8.87mL per 100g body weight. Based on this, we removed approximately 25% of air in the swim bladder using a hypodermic needle from 7 fish. We also attempted to insert air into the swim bladder. However, the injected air would most often escape the over pressurized bladder through the insertion site and float freely through the body cavity. Only one fish retained the air in the swim bladder and not in the body cavity. Six control fish were left untreated. Once treated, the fish were preserved in 50% EtOH over the course of 2 days.

The fish were μCT scanned (Fig. 1A) to generate 3D models from which COM and COB were calculated. μCT scans are most accurate when the thickness of the specimen the X-ray must penetrate does not change as the specimen rotates in the scanner. To minimize changes in thickness due to rotation, the fish were placed in the scanner vertically with their heads pointing down, with a few mL of ethanol to prevent the fish from drying during the scan. Fish were scanned (Fig. 1a) no later than one week following fixation using a SkyScan 1275 μCT scanner (Bruker, Kontich, Belgium) at a resolution of 0.048.3mm with an x-ray source voltage of 45 kV and current of 222 mA. Projection images were generated over 360° with a 0.1° rotation step and 6 averaging frames. During reconstruction of the scans, thresholding, ring artifact reduction, and beam hardening correction values were consistent across all scans using NRecon (Bruker, Kontich, Belgium). We measured the horizontal (X) distance as the distance from the tip of the mandible and the vertical (Z) location as the distance from the location where the pelvic fins meet the body. To test any effects of orientation on COM/COB locations, we additionally scanned four fish in a vertical head-up orientation.

**Figure 1.**
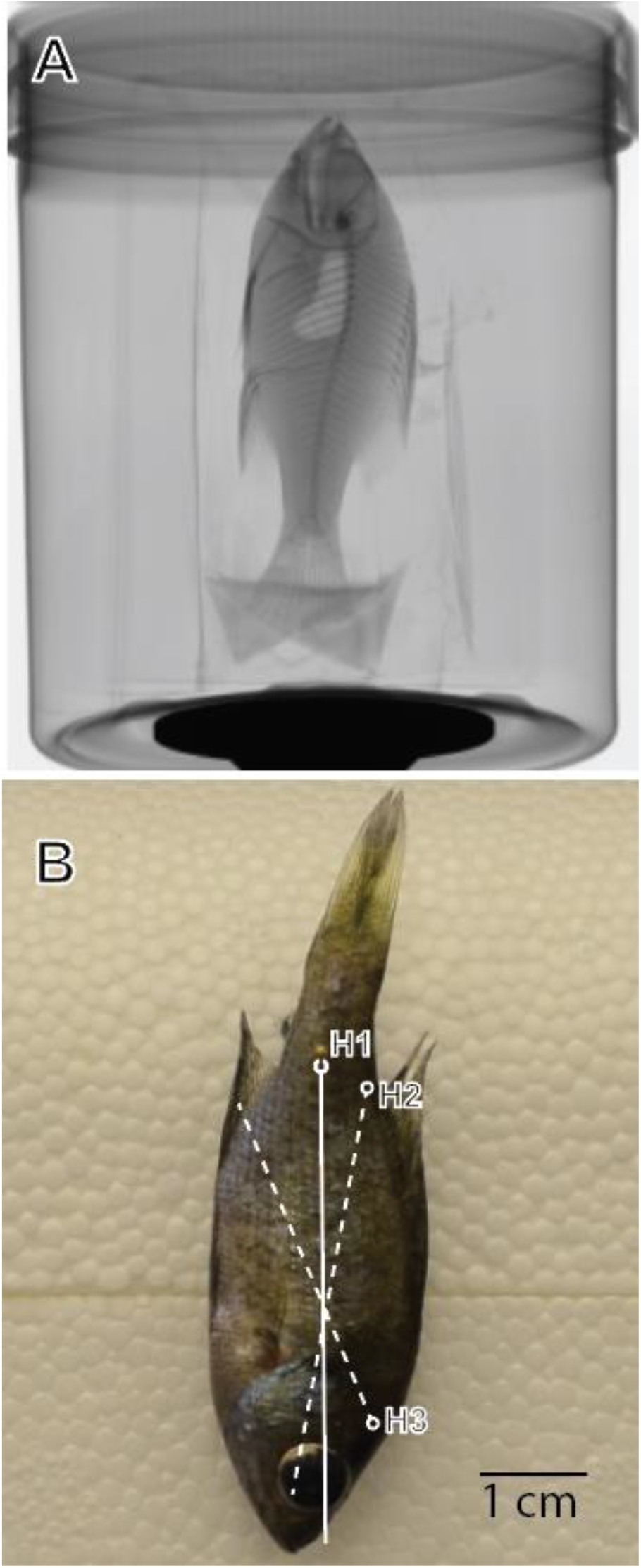
A. Fish were μCT scanned to calculate the center of mass and center of buoyancy. The panel shows a 2D planar X-ray image of a fish, taken just before scanning. B. Measurement of the center of mass via the plumb line technique. Shown here is an image of a fish hanging from suspension point H1. A line perpendicular to the gravitational vector that goes through H1 is the solid white line. H2, H3, and their associated dotted lines represent the two other suspension points and their associated vertical lines and have been overlayed on the H1 image. The COM is located at the intersection of these lines.

### Density calibration

We used the intensity at each voxel as a measure of density. Voxel intensity is a measurement of x-ray absorption, also called the attenuation coefficient (mm^-1^). The amount of x-ray absorption at a given voxel is determined by density, but it is also influenced by chemical composition at that point (Du Plessis et al., 2013). For homogeneous objects, digital determination of COM location is more accurate and shows less variability between estimates than the plumb line technique (Macaulay et al., 2017). While this is true for an object of uniform composition, voxel intensity does not necessarily scale linearly across tissue types. To account for this, recent studies have used calibrated μCT scans to measure the location of the COM (Durston et al., 2022). Calibrated scans use tissue characterization samples with a known density (also called phantoms) to establish a correlation between voxel intensity and density.

We estimated a scaling relationship for hard tissues (bones and scales) and a separate scaling factor for soft tissues (muscle, viscera, fat, etc.). To create a scaling relationship for hard tissue, we included a 5 mm phantom with cylindrical inserts containing four known densities of calcium hydroxyapatite in two scans (QRM, Möhrendorf, Germany). We isolated each sample in the phantom and recorded the average voxel intensity and compared it to the density of the sample. To create a scaling relationship for soft tissues, using 3D Slicer (Version 4.11, Fedorov et al., 2012) we measured the voxel intensity and density of two easily differentiated tissues with different gray values: anterior dorsal epaxial muscle and the lens of the eye. For each scan, we recorded the voxel intensity of the dorsal epaxial muscle and one of the eye lenses, then calculated the average value for each tissue. We calculated the density of these tissues from dissected fish,which could not be scanned, because the scanning process dries out the tissue over time. To calculate the density we measured the mass of the tissue sample and divided by its volume measured by the weight of water displacement per sample.

### COM and COB location calculation

We used the reconstructed virtual fish models to calculate the location of the COM and COB. If there was any bending of the fish in the scan, the fish was digitally straightened using unwind.exe (Williams et al., 2020). This process also guaranteed that the fish were oriented along the same X,Y,Z coordinate system. We then used 3D Slicer (Version 4.11, Fedorov et al., 2012) to isolate only the voxels that comprised the fish, removing any voxels including support material or background.

Custom Matlab software (R2020b; Mathworks, Inc, Natick, MA) was used to calculate the COM and COB from the isolated fish reconstruction. The COB was calculated by determining the geometric center of the isolated fish volume:

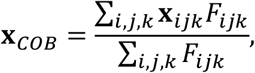

where **x**_*ijk*_ = [*x*_*i*_ *y*_*j*_ *z*_*k*_] is the 3D location of the voxel with indices *i, j*, and *k*, and *F*_*ijk*_ is 1 for voxels inside the fish’s volume and 0 otherwise.

The COM was calculated as the average location in the body, weighted by density:

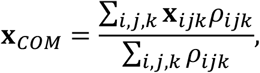

where *ρ*_*ijk*_ is the density of the voxel at indices *i, j*, and *k*. Different scaling relationships were used to determine the density of each voxel for hard and soft tissue (see “Density calibration” above).

We measured the location of the COM of 4 fish using the plumb line technique (also called the suspension method; Fig. 1B). The plumb line technique locates an object’s COM by that object and a weighted string (plumb line) from the same point of rotation. The object will rotate until it’s COM is below the point of rotation and the plumb line is used to draw a line on the object. The object is then suspended from a new point of rotation and the plumb line is used to draw a second line. The intersection of those lines indicates the objects COM. We pierced each fish with a dissecting pin and hanged the sample so that it could rotate freely. We photographed the fish in a lateral view. A weighted plumb line was suspended and included in the image to indicate the gravity vector. This process was repeated two more times from two different points of rotation, a total of three times for each fish. We used Photoshop (2021, Adobe, Venture, CA) to draw a vertical line through the suspension point parallel to the plumb line. We repeated drawing process for each imaged suspension point. The three images were digitally rotated so that the fish was aligned (Fig. 1B). The three lines drawn from the hanging point on each image intersected to create a triangle, and the centroid of this triangle was taken as the center of mass.

### Swim bladder centroid determination

We calculated the location of the centroid of the swim bladder to compare its relative location in fish in the head-up and head-down orientations. This analysis included two fish which were not used elsewhere in the analysis: these individuals contained air bubbles in the mouth or gut which would affect COM calculations and were therefore excluded from other analyses. However, these bubbles would not affect the shape of the swim bladder, so they were included in the swim bladder centroid analysis.

### Statistical analysis

All measured values are reported as mean ± standard deviation. We used a one-sample t-test to measure if the difference between the measured mass of the fish and the estimated mass of the fish was different than zero. We estimated regressions for the density and mass relationship as well as the swim bladder and COM/COB location relationships with a least squares regression.

## Results

### Density estimation

We calculated a scaling relationship between density and voxel intensity for hard and soft tissue. These values yielded the scaling relationships (Fig. 2A). To determine if these relationships were accurate we added up the estimated density of each voxel to estimate the mass of the fish (Fig. 2B). Using scaling relationships for hard and soft tissues we found that voxel intensity could be used to accurately estimate the mass of the fish (r^2^ = 0.9971). The difference in the estimated and true mass was not significantly different from zero (t = 1.45, p = 0.17), indicating a 1:1 relationship between the actual mass and estimated mass.

**Figure 2.**
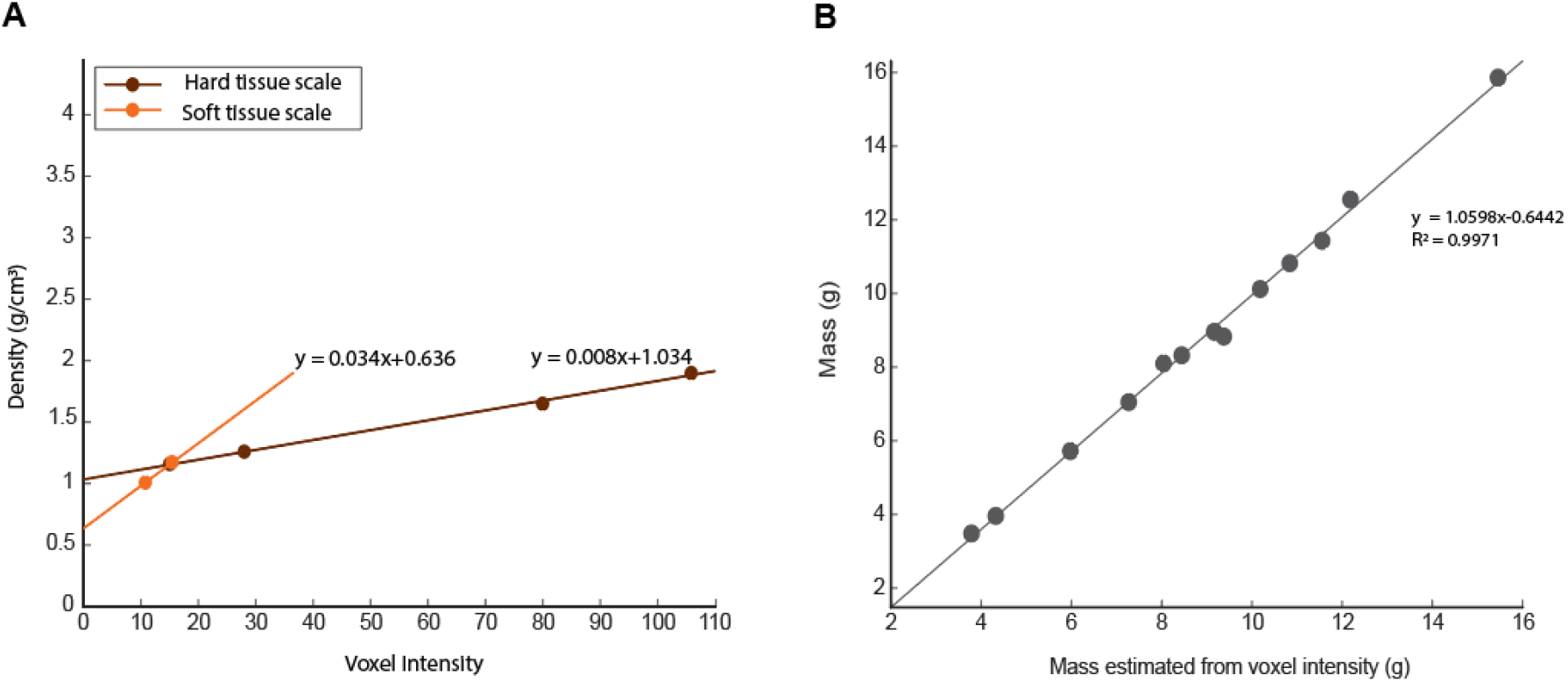
A. The scaling relationships for voxel intensity and density for hard tissue (brown) and soft tissue (orange). B. The mass of a fish vs the mass of the same fish estimated from voxel intensity.

We measured the location of the COM using both the digital and hanging techniques in four individuals. For repeated measurements taken from the same individual, the measured COM location varied by 2.35±2.9 % BL. The COM locations calculated from the μCT scan reconstructions were 2.59±3.18 % BL from the COM locations determined by hanging on the same fish, within the range of precision of the plumb line technique.

### The swim bladder and the location of the COM and COB

The bluegill individuals we measured had a mean mass of 9.3 ± 3.49 g (n = 15) and mean length of 8.38 ± 0.01 cm.

In most fish, the COM was located posterior and ventral to the COB (Fig. 3). On average, the COM was located 0.442 ± 0.008 BL from the snout and 0.193± 0.01 BL from the ventral surface, and the COB is located 0.441 ± 0.0072 BL from the snout and 0.194±0.01 BL from the ventral surface (Fig. 3A). Comparing the location of the COM relative to the COB COB, we found that there is always some distance between the COM and COB (Fig. 3B). For fish scanned head-up in the scanner, the COM is 0.0012±0.0023 BL posterior and 0.00045±0.00051 BL ventral to the COB on average. This arrangement would create a small head-up pitching torque, which was 2.04 μNm ± 2 μNm.

**Figure 3.**
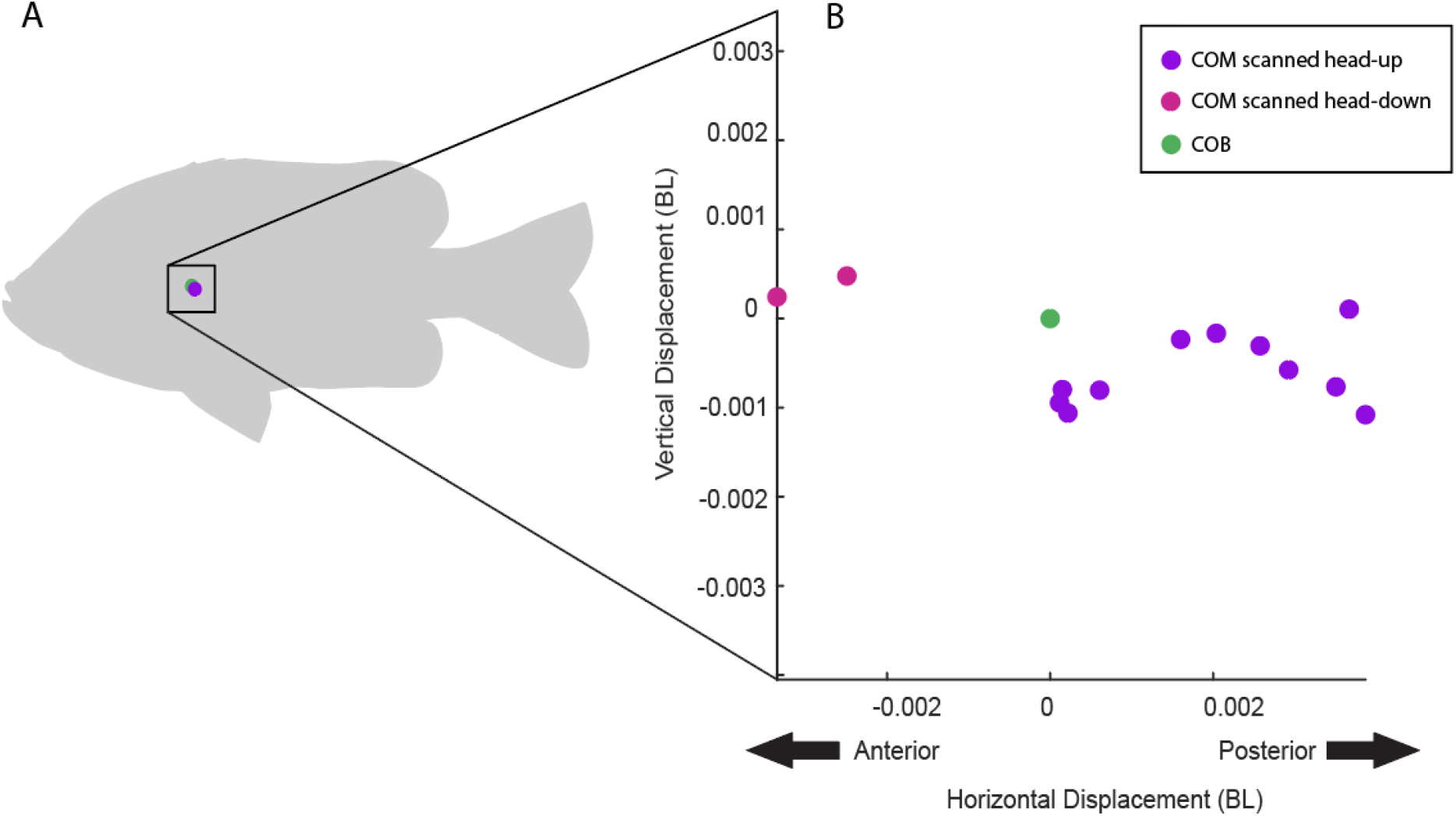
The COM is usually located posterior and ventral to the COB. A. The average location of the COM and COB. B. Each COM measurement relative to the COB.

We found no correlation between the volume of the swim bladder and the location of the COM within the fish (p = 0.82, R^2^ = 0.0046, Fig. 3).

Fig. 4 shows the relative locations of the COM and COB in fish with swim bladder volumes ranging from 1.9 to 7.6% total fish body volume. In untreated fish, the mean swim bladder size is 6.2%± 1.2% of the total body volume (range from 4% to 7.6% body volume). We were able to remove air from 7 individuals. The mean swim bladder volume for fish with air removed was 4.6 ± 1.1% total fish body volume. The swim bladder was 7.2% total fish body volume in the individual with air added to the bladder.

**Figure 4.**
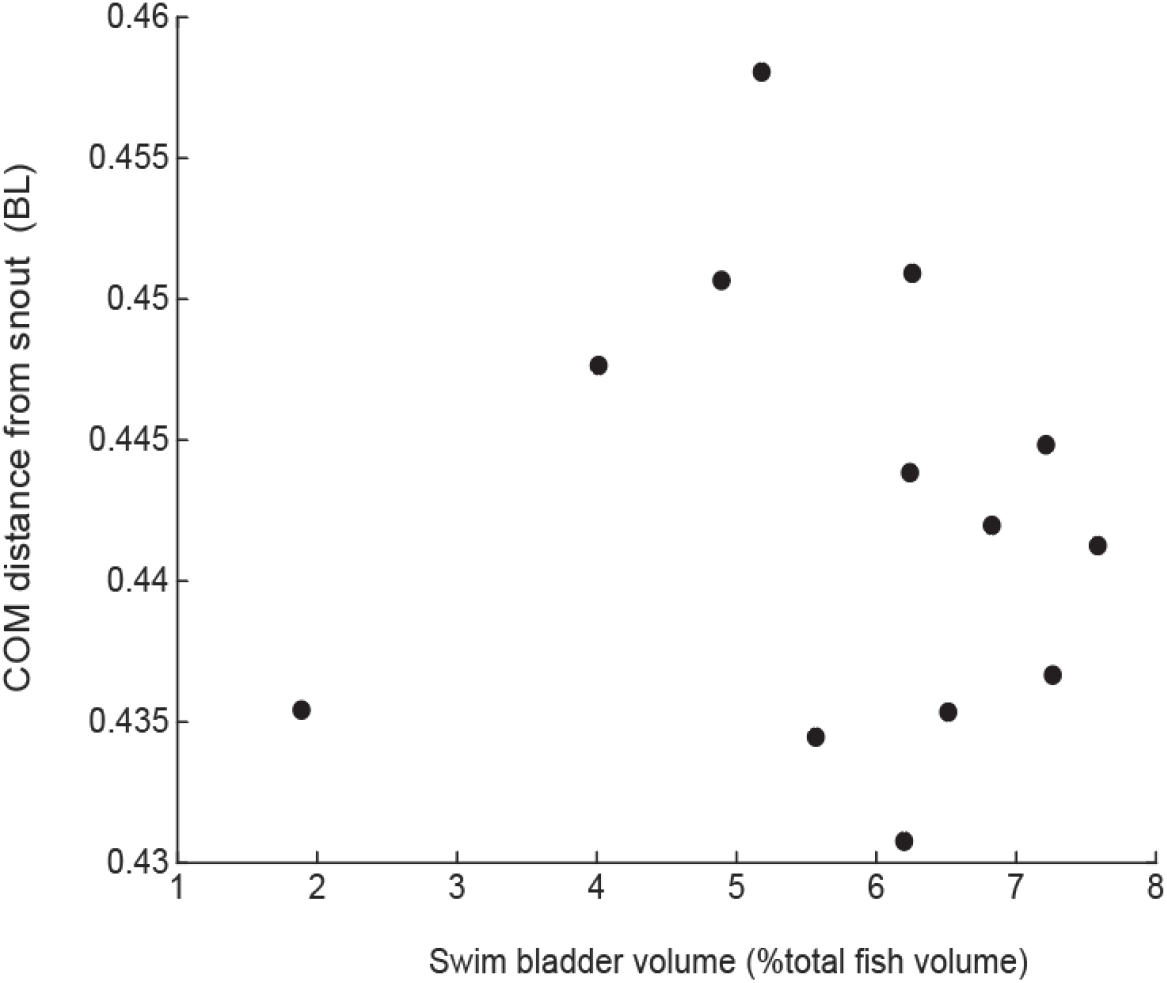
The center of mass location does not depend on the swim bladder volume. The plot shows the horizontal location of the center of mass compared to the volume of the swim bladder.

We also found no significant relationship between swim bladder volume and the total distance between the COM and COB (Fig. 5) (p = 0.74, R^2^ = 0.0102). Further, we did we find any correlation between the swim bladder volume and the horizontal (p=0.83, R^2^ = 0.0046) or vertical (p=0.89, R^2^ = 0.001) components of the COM/COB displacement.

**Figure 5.**
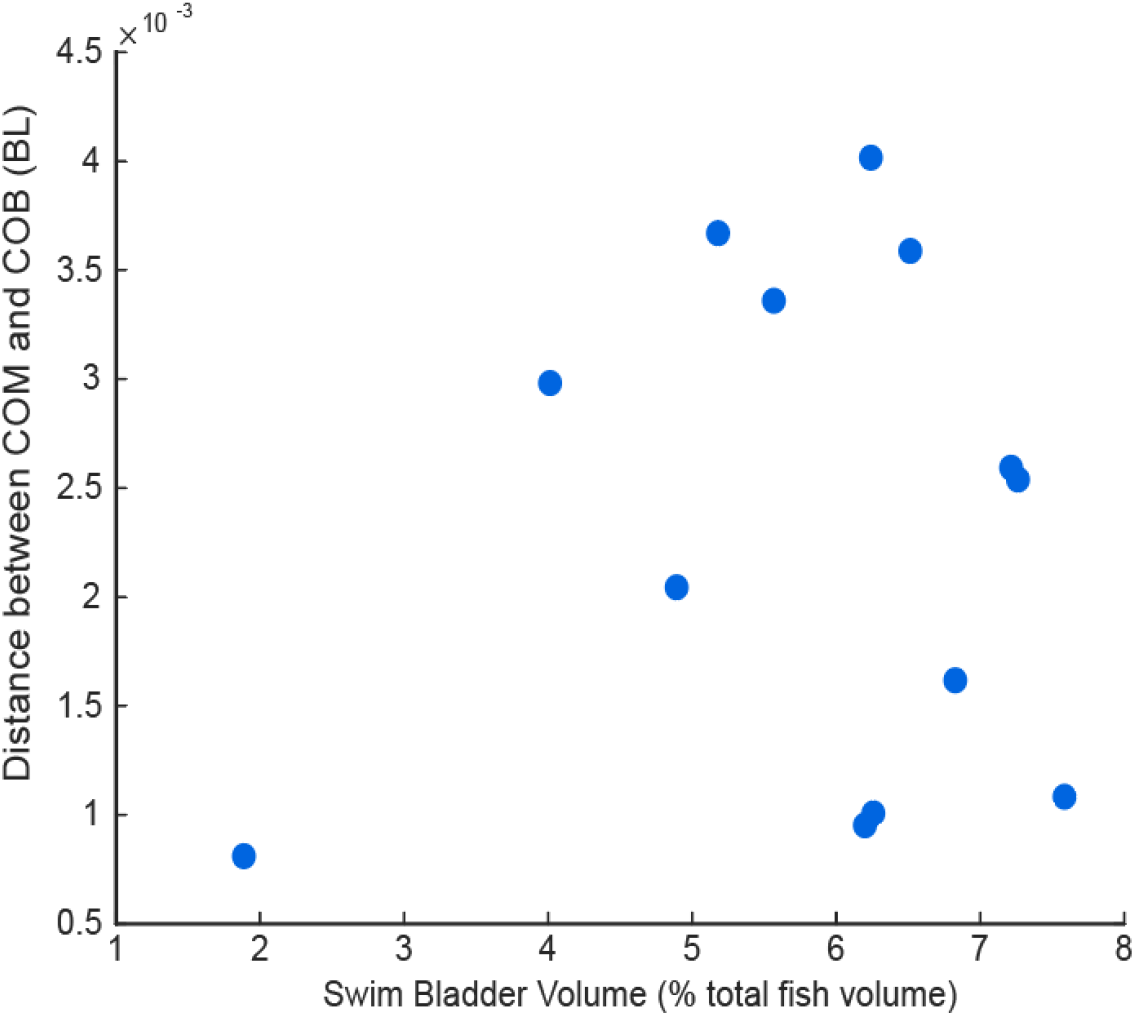
The distance between the COM and COB does not depend on swim bladder size.

Although the swim bladder volume does not affect the location of COM or COB, the swim bladder location does have an effect. In fish scanned head up, the swim bladder is positioned significantly (p =<0.001) more posteriorly than in fish scanned head-down (Fig. 6). On average, the centroid of the swim bladder in fish scanned head-down (n=4) is located 0.53±0.015 BL from the snout. The centroid of the swim bladder in fish scanned head-up (n=11) is located 0.45±0.0097 BL from the snout. In all fish, the swim bladder is just ventral to the vertebrae. In fish scanned head up, the bladder extends from posterior to the neurocranium and terminates before the first haemal/interhaemal arches. In fish scanned head-down, there is a larger distance between the posterior edge of the cranium and the bladder and the bladder straddles the first haemal/interhaemal arches.

**Figure 6.**
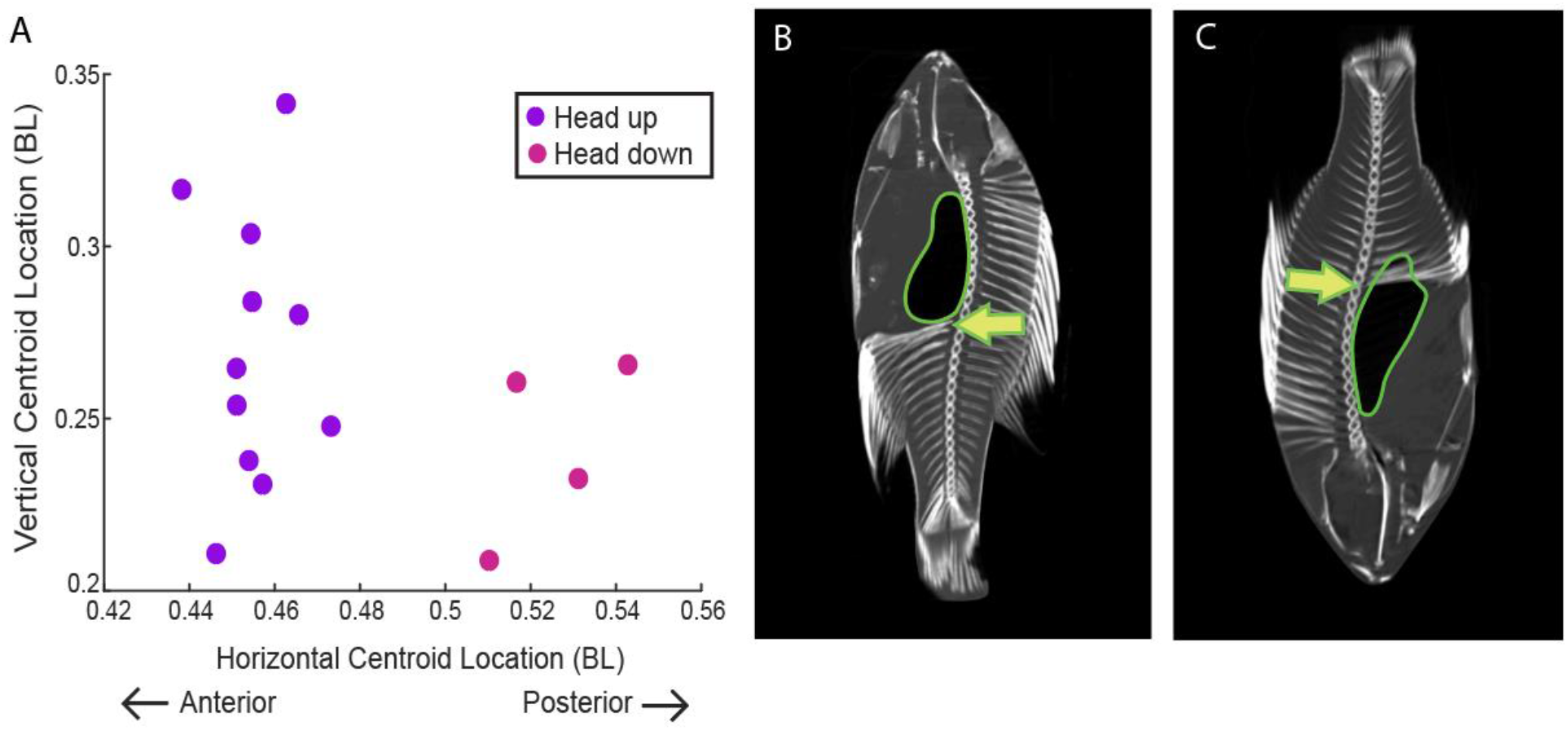
The location of the centroid of the swim bladder in bluegill sunfish depends on the fish’s orientation in the scanner. A. The centroid location in fish scanned head up (purple) and head down (pink). B. A fish placed in the scanner head up. C. A fish placed in the scanner head down. The swim bladder is outlined in green and the level of the first haemal arch is indicated with an arrow.

## Discussion

The swim bladder can change static stability in fishes. Straightforwardly, changes in swim bladder volume change the magnitude of the buoyant force. However, the degree to which that force contributes to potentially destabilizing torques depends on the relative locations of the COM and COB. On average, we found that the COM is located posterior and ventral to the COB in bluegill sunfish when they were scanned in a head-up orientation. These locations of the COM and COB generate a head-up pitching torque. Conversely, fish placed in the scanner head-down had a COM located anterior and dorsal to the COB, a configuration which would result in a head-down pitching torque. This change in COM/COB positions result from the change in shape and placement of the swim bladder at different pitching angles, because the centroid of the swim bladder is located further posterior in fish in the head-down orientation when compared to the head-up orientation (Fig. 5).

While we found no correlation between swim bladder volume and COM/COB positions, we did find that the redistribution of tissue from one pitching extreme to the other drives a change in the relative locations of the COM and COB, resulting in a change in torque direction.

### Destabilizing pitching torques can be easily countered by bluegill

The horizontal distance between the COM and COB was small, on average 0.0012 BL. Using the horizontal distance as the length of the moment arm, we calculated that the fish experience an average pitching torque of 2.04 μNm. To estimate the ability of the bluegill to counter this torque we used the highest possible torque from our observations. The largest distance between the COM and COB was 0.75 mm and the largest buoyant force was 0.081 N. Assuming the moment arm is perfectly perpendicular to the pull of gravity, the largest possible destabilizing torque is 62.9 μNm, which is 20 times greater than the average. Drucker and Lauder (1999) estimated that bluegill can produce up to 24 mN of horizontal force with the pectoral fins on the upstroke. If we assume that the center of pressure of the fin, where the force is centered, is located approximately 1.2cm from the center of mass, then the fin can generate a torque of 288 μNm, much more than the torque resulting from the buoyant force. Bluegill can therefore easily produce more torque than needed to counteract even the maximum passive torque from buoyancy.

### Changes in swim bladder volume do not change the static stability orientation

Contrary to our hypothesis, we found no correlation between swim bladder size and the distance between the COM and COB, comparing swim bladders that varied in size over a range of 5.7% total body volume. This suggests that fish do not change in static stability as they change depth.

Jones (1951) and Alexander (1959d) found that the swim bladder of many fish changes volume with pressure in close accordance with Boyle’s law (P_1_V_1_ =P_2_V_2_, where P is pressure and V is volume). Using Boyle’s law, we can estimate that a decrease in 5.7% swim bladder volume is equivalent to a bluegill swimming from a depth of 1 m down to about 3.8 m in fresh water. Bluegill can be found in lakes and ponds deeper than 4 m so it is possible that they might experience a greater change in swim bladder size than we measured. However, a fish that seeks to maintain a constant buoyant force must maintain a constant swim bladder volume, regardless of depth. Moreau (1876) demonstrated that, under persistent pressure changes, fish actively change the gas content in swim bladder such that volume of the bladder is constant despite hydrostatic pressure. Any changes in swim bladder volume resulting from decreased or increased water pressure are likely to be temporary as the fish adjusts the amount of gas in the bladder to meet the new equilibrium.

### Static stability: it’s not so static

While static stability likely does not change as fish change the volume of their swim bladders, the stability may change with body orientation. We found that the distribution of air in the body cavity changes with pitch angle. We scanned fish in two orientations: head-up and head-down. Fish scanned in either pitch extreme have opposite COM/COB configurations, and therefore, different stable orientations. In fact, the COM is below the COB in each orientation, making the fish relatively stable in each orientation. This change in the relative locations of COM and COB likely results from the redistribution of tissues. Air rises upward within the swim bladder to occupy the highest point available, while the heavier soft tissues “sink” downward to occupy lower points in the gut cavity. The centroid of the swim bladder in fish scanned head up is located 0.08 BL more anteriorly compared to the swim bladder centroid of fish scanned head-down. These differences suggest that the location of COM may be dynamic and dependent on pitch, and that there may be an intermediate pitch angle at which the COM and COB are vertically aligned, thus producing no destabilizing pitching or rolling torques. Fishes (He and Wardle 1986; Webb 1993; Wilga and Lauder 1999, 2000,2001), including bluegill (Eidietis, Dorrester, and Webb, 2002) have indeed been observed to change pitch in response to destabilizing forces. Pitching behavior may thus be an important and simple behavior fish use to minimize static instability.

### Calibrated digital COM measurement is as at least as accurate as the plumb-line technique

The extremely small distances between the center of mass and center of buoyancy have made it difficult to determine the relative location of the two points. Earlier studies have measured the location of COM and COB using different techniques such as serial sectioning, balancing, and the plumb-line (suspension) technique. These measuring techniques can introduce error. Serial sectioning results in loss of water and tissue as the fish is sliced and limits COM location determination to one axis. Traditional plumb line techniques require the creation of a physical model to locate the COB. Slight differences in 3D orientation of the model or fish while measuring can lead to differing location measurements. These limitations are not trivial. Where it is reported, repeated measurements of the location of the COM and COB locations in fishes varied by ± 2% body length, which is greater than the distance between the two points (Weihs 2011).

Studies reporting on homogeneous objects have shown that using a μCT scan to digitally locate the COM of the object can be more accurate than using the plumb line method (Macaulay et al., 2017). CT or MRI scans have been used to find the COM in humans (Pearsall et al., 1996), dogs (Amit et al., 2009), and birds (Macaulay et al., 2017). However, μCT scans also have limitations, because animals are not made of homogeneous materials. Using voxel intensity from μCT scans to calculate the center of mass is one potential source of error because intensity is not a direct measure of density, but actually a measure of x-ray absorption, which can vary across tissue types. Our study, like others (Durston et al., 2022), accounted for this by calibrating scans with samples of specific tissues of known densities.

We used separate scaling factors for hard and soft tissues to calibrate our scans and account for non-density-related changes in x-ray absorption. Using this calibration, we were able to estimate the body mass within 2.4% ±3.86% from the reconstructed scans. We also compared the location of the COM determined from calibrated μCT scan to the plumb line method. We found that on average, our digitally determined location was 2.6% BL from the average measured plumb line COM location. This is comparable to the variability from repeated measures of ± 2% body length reported by Weihs (2011). We conclude that using calibrated μCT scans to calculate the location of the center of mass for a fish is at least as accurate as traditional techniques.

Using μCT scans has an additional advantage: they provide a precise reconstruction of fish volume and body shape, so they can be used to estimate COB location with extreme accuracy. Use of the plumb line technique requires the creation of a separate animal model made from a homogeneous material (Alexander (1959d) used wood, Web and Weihs (1994) used plaster) to determine the center of buoyancy. Then researchers would compare measurements between the model and the actual animal. The CT-based method uses the same digital model to calculate both COM and COB, thus removing error introduced by comparing the fish to a separate model.

## Conclusion

Bluegill sunfish are very close to being neutrally stable, but the displacement between the COM and COB does generate small destabilizing pitching torques. Changes in swim bladder volume are not correlated with changes in the distance between the COM and COB. However, the relative locations of these two points can change depending on the pitching orientation of the fish. This change may be driven by the redistribution of swim bladder and soft tissues as the fish changes pitch angle. Active changes in pitching angle may be a strategy a fish can use to minimize instabilities resulting from the interaction of the gravitational and buoyant forces.

## Data Availability

All data will be made available via LabArchives with a citable DOI.

## References

Alexander, R. McN. (1959d), The physical properties of the swimbladders of fish other than Cypriniformes, J. exp. Biol. 36, 347–355.

Aleyev, Y.G. 1977. Nekton, Dr. W. Junk bv Publishers, Dordrecht, the Netherlands

Amit, T., Gomberg, B. R., Milgram, J., & Shahar, R. (2009). Segmental inertial properties in dogs determined by magnetic resonance imaging. Veterinary Journal, 182(1), 94–99. https://doi.org/10.1016/j.tvjl.2008.05.024

Du Plessis, A, Meincken, M, & Seifert, T. (2013). Quantitative determination of density and mass of polymeric materials using microfocus computed tomography. Journal of Nondestructive Evaluation, 32(4), 413–417.

Durston, N. E., Mahadik, Y., & Windsor, S. P. (2022). Quantifying avian inertial properties using calibrated computed tomography. Journal of Experimental Biology, 225(1). https://doi.org/10.1242/JEB.242280

Fedorov, A., Beichel, R., Kalpathy-Cramer, J., Finet, J., Fillion-Robin, J. C., Pujol, S., Bauer, C., Jennings, D., Fennessy, F., Sonka, M., Buatti, J., Aylward, S., Miller, J. v., Pieper, S., & Kikinis, R. (2012). 3D Slicer as an image computing platform for the Quantitative Imaging Network. Magnetic Resonance Imaging, 30(9), 1323–1341. https://doi.org/10.1016/j.mri.2012.05.001

Jones, F. R. H. (1951). The swimbladder and the vertical movement of teleostean fishes. I. Physical factors. J. exp. Biol. 28, 553–66.

Macaulay, S., Hutchinson, J. R., & Bates, K. T. (2017). A quantitative evaluation of physical and digital approaches to centre of mass estimation. Journal of Anatomy, 231(5), 758–775. https://doi.org/10.1111/joa.12667

Mcnabb, R. A., & Mecham, J. A. (1971). The effects of different acclimation temperatures on gas secretion in the swimbladder of the bluegill sunfish, Lepomis macrochirus. Comparative Biochemistry and Physiology -- Part A: Physiology, 40(3), 609–616. https://doi.org/10.1016/0300-9629(71)90245-3

Pearsall, D. J., Reidt, J. G., & Livingston~, L. A. (1996). Segmental Inertial Parameters of the Human Trunk as Determined from Computed Tomography. In Annals of Biomedical Engineering (Vol. 24).

Smits, A. J. (2000). A Physical Introduction to Fluid Mechanics.

Steen, J. B. (1970). The swim bladder as a hydrostatic organ. Fish Physiology, 4(C), 413–443. https://doi.org/10.1016/S1546-5098(08)60135-1

Walker, J. (2015) Fundamentals of Physics Vol 1.

Webb, P. W., & Weihs, D. (1994). Hydrostatic stability of fish with swim bladders: not all fish are unstable. Canadian Journal of Zoology, 72(6), 1149–1154. https://doi.org/10.1139/z94-153

Webb, P. W., & Weihs, D. (2015). Stability versus Maneuvering: Challenges for Stability during Swimming by Fishes. Integrative and Comparative Biology, 55(4), 753–764. https://doi.org/10.1093/icb/icv053

